# Protein language model-based end-to-end type II polyketide prediction without sequence alignment

**DOI:** 10.1101/2023.04.18.537339

**Authors:** Jiaquan Huang, Qiandi Gao, Ying Tang, Yaxin Wu, Heqian Zhang, Zhiwei Qin

## Abstract

Natural products are important sources for drug development, and the precise prediction of their structures assembled by modular proteins is an area of great interest. In this study, we introduce DeepT2, an end-to-end, cost-effective, and accurate machine learning platform to accelerate the identification of type II polyketides (T2PKs), which represent a significant portion of the natural product world. Our algorithm is based on advanced natural language processing models and utilizes the core biosynthetic enzyme, chain length factor (CLF or KS_β_), as computing inputs. The process involves sequence embedding, data labeling, classifier development, and novelty detection, which enable precise classification and prediction directly from KS_β_ without sequence alignments. Combined with metagenomics and metabolomics, we evaluated the ability of DeepT2 and found this model could easily detect and classify KS_β_ either as a single sequence or a mixture of bacterial genomes, and subsequently identify the corresponding T2PKs in a labeled categorized class or as novel. Our work highlights deep learning as a promising framework for genome mining and therefore provides a meaningful platform for discovering medically important natural products.

## Introduction

Bacterial type II polyketides (T2PKs) are valuable natural products with potent biological activities and comprise a family of structurally related molecules (*1, 2*). Illustrative examples include tetracycline, doxorubicin and plicamycin. They are primarily biosynthesized by type II polyketide synthases (T2 PKSs), which catalyze the formation of carbon skeletons in an ordered manner (*3*). The biosynthetic system is characterized by a minimal set of gene products, of which the most crucial enzymes are the monofunctional heterodimeric β-ketosynthase pair KS_α_/KS_β_, which catalyze the iterative Claisen condensation using acetyl- and malonyl-CoA as building blocks for chain elongation and determine the chain length and overall topology (Fig. 1A), respectively. In addition, a malonyl transacylase (MT) and an acyl carrier protein (ACP) were taken together with KS_α_/KS_β_ to constitute the minimal T2 PKS systems (*4*). The core skeleton of T2PKs is highly correlated with the KS_α_/KS_β_ protein structure. Hillenmeyer et al. observed correlations between KS_β_ protein phylogeny and the building blocks of T2PK skeletons (*5*), while Chen et al. utilized KS_β_ as a biomarker to construct a coevolutionary statistical model (phylogenetic tree) to expand the T2PK biosynthetic landscape (*6*). However, the above and other (*7-9*) methods frequently rely on multiple sequence alignments, which are time-consuming and do not effectively represent protein structural information (*10, 11*).

**Fig. 1.**
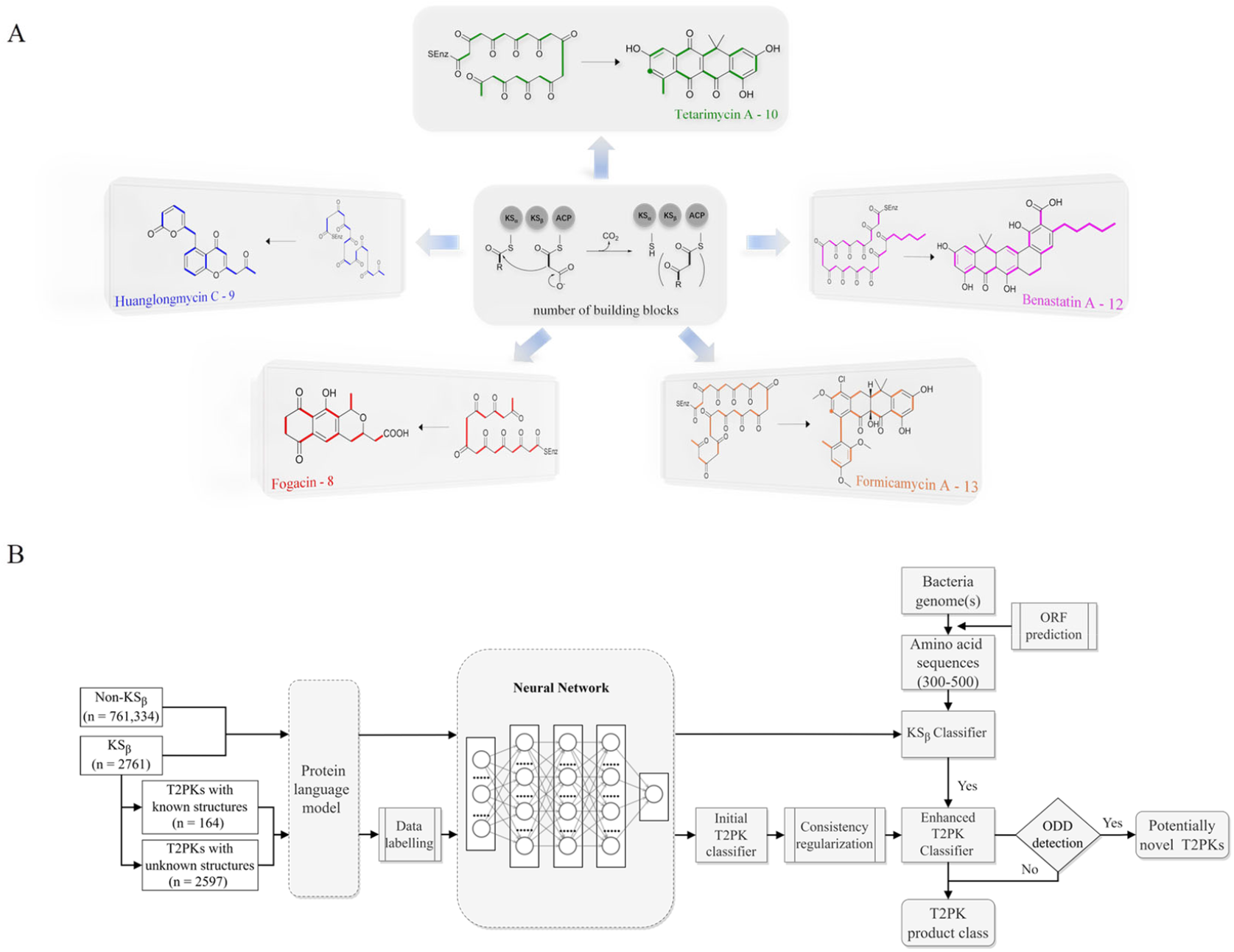
Illustration of the machine learning-based algorithm employed in this study. (**A**) Depiction of various type II polyketide natural products (T2PKs) and their corresponding poly-β-ketone intermediates during the biosynthesis process. The bold bonds in different colors indicate the number of building blocks incorporated into the polyketide backbone (intermediate chain length), such as 8 (octaketide, fogacin, red), 9 (nonaketide, huanglongmycin C, blue), 10 (decaketide, tetracycline, green), 12 (dodecaketide, benastatin, purple), and 13 (tridecaketide, formicamycin, orange), respectively. (**B**) Workflow of DeepT2. The training dataset comprised KS_β_ and non-KS_β_ sequences. To convert the protein sequences into embeddings that represent protein structural information, various PLMs including SeqVec, ESM-1b, ESM-2, ProtT5-XL-U50 and ProtBert-BFD were employed initially; KS_β_ classifier and T2PK classifier are established and trained by embeddings.

Several artificial intelligence (AI)-based natural product discovery models have been proposed in recent years due to rapid data accumulation and digital transformation as well as the accelerated development of AI technology (*12-14*). Among these frameworks, deep learning has demonstrated exceptional performance in classification tasks, specifically in the area of distinguishing new and unseen data (*15, 16*). For instance, DeepBGC, trained in a deep learning framework, exhibited high efficiency and large scale in predicting the natural product class (*17*). More recently, protein language models (PLMs) based on self-supervised learning have shown remarkable ability to convert individual protein sequences into embeddings that describe the homology between multiple protein sequences and potentially capture physicochemical information not encoded by the existing methods (*18-20*). The application of general PLMs to convert sequences into embeddings, which serve as inputs for deep learning models, effectively overcomes the few-shot learning challenge for specific biomolecular property predictions (*21, 22*). In addition, leveraging the large amount of unlabeled data available through a semisupervised framework can further improve model performance (*23, 24*). These advancements inspired us to move forward in understanding T2PKS with PLM, training a robust model with unexplored sequences stored in metagenome data, and eventually finding an effective linker to connect their biosynthetic enzymes and probable chemical structures.

To gain a better approach for the discovery of T2PKs, an end-to-end model named DeepT2 was developed in this study. This model employs multiple classifiers to expedite the translation from protein sequences to the T2PK product class and identify any potential new compounds beyond the established groups. Notably, the model is free of sequence alignment and comprises four main components: (i) sequence embedding: the protein sequences were converted into vector embeddings using a pretrained PLM called EMS-2 (*20*); (ii) data labeling: the KS_β_ dataset with known corresponding chemical structures was initially split into five classes for labeling based on the total number of biosynthetic building blocks, which was later reclassified into nine classes through dimension reduction and clustering processes; (iii) classifier development: this was used for both KS_β_ sequences and T2PKs classification; and (iv) novelty detection: Mahalanobis distance-based novelty detection (*25*) was applied to identify any potential new compounds beyond the nine established groups. Remarkably, we leveraged DeepT2 to detect KS_β_ from microbial genomes and successfully identified four T2PKs as categorized in our classifiers. This work paves a promising avenue to further explore the potential of the existing reservoirs of T2PK biosynthetic gene clusters (BGCs) and therefore expand the chemical space of this medically important natural product family.

## Results

### DeepT2 model architecture

As shown in Fig. 1, the purpose of this study was to develop a methodology for predicting the T2PK class using KS_β_ sequences from bacterial genomes as input. To achieve this, we used an ensemble of multiple classifiers to determine whether a given protein sequence belongs to KS_β_, assessed whether it fell within or outside the existing labeling product classes and eventually predicted the product class. General protein language models, including SeqVec, ESM-1b, ESM-2, ProtT5-XL-U50 and ProtBert-BFD, which were embedded with structural features from large-scale protein sequence datasets, were employed to maximize the use of limited labeled, unlabeled, and non-KS_β_ sequences by converting them into embeddings containing structural representations using the idea of transfer learning. In this work, the terms of ‘labeled KS_β_’ and ‘unlabeled KS_β_’ indicate whether their corresponding chemical structures are known. KS_β_ and T2PK classifiers were trained by the datasets from the specific protein sequences and chemical structures. To construct a robust T2PK classifier, we applied supervised UMAP (uniform manifold approximation and projection) and HDBCAN (hierarchical density-based spatial clustering of applications with noise) on the KS_β_ embedding to generate more appropriate class labels and trained the model with unlabeled data on the basis of consistency regularization. Furthermore, the Mahalanobis distance-based algorithm was applied on each feature layer of the T2PK classifier to perform novelty detection and avoid the problem of overconfident Softmax-based classifiers. The methodology enabled us not only to classify the known group of T2PKs but also to detect potential novel classes of T2PKs from unknown KS_β_ protein sequences. Detailed results of each process are presented below.

### Development of the KS_β_ classifier

We obtained a collection of 164 labeled KS_β_ sequences with known corresponding natural product structures (Table S1) as well as an additional set of 2597 unlabeled KS_β_ sequences sourced from the RefSeq database, whose associated natural product structures remain unknown (Table S2). A total of 2761 (164 + 2597) KS_β_ sequences and 761,334 non-KS_β_ sequences were then split into training, validation, and test datasets, as described in the materials and methods section. Prior to constructing the classifier, we employed five general PLMs, SeqVec, ESM-1b, ESM-2, ProtT5-XL-U50 and ProtBert-BFD, to vectorize each protein sequence for embedding and we observed that the learned representations of ESM-2 and ProtT5-XL-U50 exhibited the best performance compared with the others in distinguishing KS_β_ from non-KS_β_ sequences, as revealed by the results of dimension reduction (Fig. S1). In this study, we favored ESM-2 because it has the largest parameter size (over 3 billion) and has better performance on the T2PK classifier (see lateral session). Next, we trained the KS_β_ embeddings obtained by the PLMs using four machine learning algorithms, including random forest, XGBoost, support vector machine (SVM) and multilayer perceptron (MLP), and found that MLP and SVM achieved the best results, with an AUROC of 1 and an F1 score of 1 in the classification on the test dataset (Table S3).

### Relabeling 164 T2PKs with constrained optimization approach

Data labeling is a crucial part of data preprocessing for machine learning, especially for supervised learning (*26*). This is also the most challenging task in this work. Based on prior knowledge, 164 KS_β_ embeddings with known chemical structures were categorized into five groups according to the building block number of their corresponding to T2PK main skeleton, namely, 8, 9, 10, 12 and 13 (Table S1). However, despite applying ESM-2 to the 164 KS_β_ sequences, imprecise representation of the distribution pattern of the 5 label embeddings was observed (Fig. S2A), probably due to the inadequate fittings between some of the class labels and the protein embeddings. To address this issue, a constrained optimization approach was developed to correct the improper class labels for each sample and further separate their local features from global features to generate new class labels (Fig. 2B). Specifically, we employed supervised UMAP and HDBSCAN as regularization techniques to constrain the latent distribution of KS_β_ embeddings and automatically tune UMAP and HDBSCAN hyperparameters using a labeling cost function to assign more appropriate class labels to embeddings. Of note, the supervised UMAP opted for this study incorporates compound skeleton labels into the optimization process to ensure that the resulting reduced-dimensional space consistently captures the features of the compound skeleton.

**Fig. 2.**
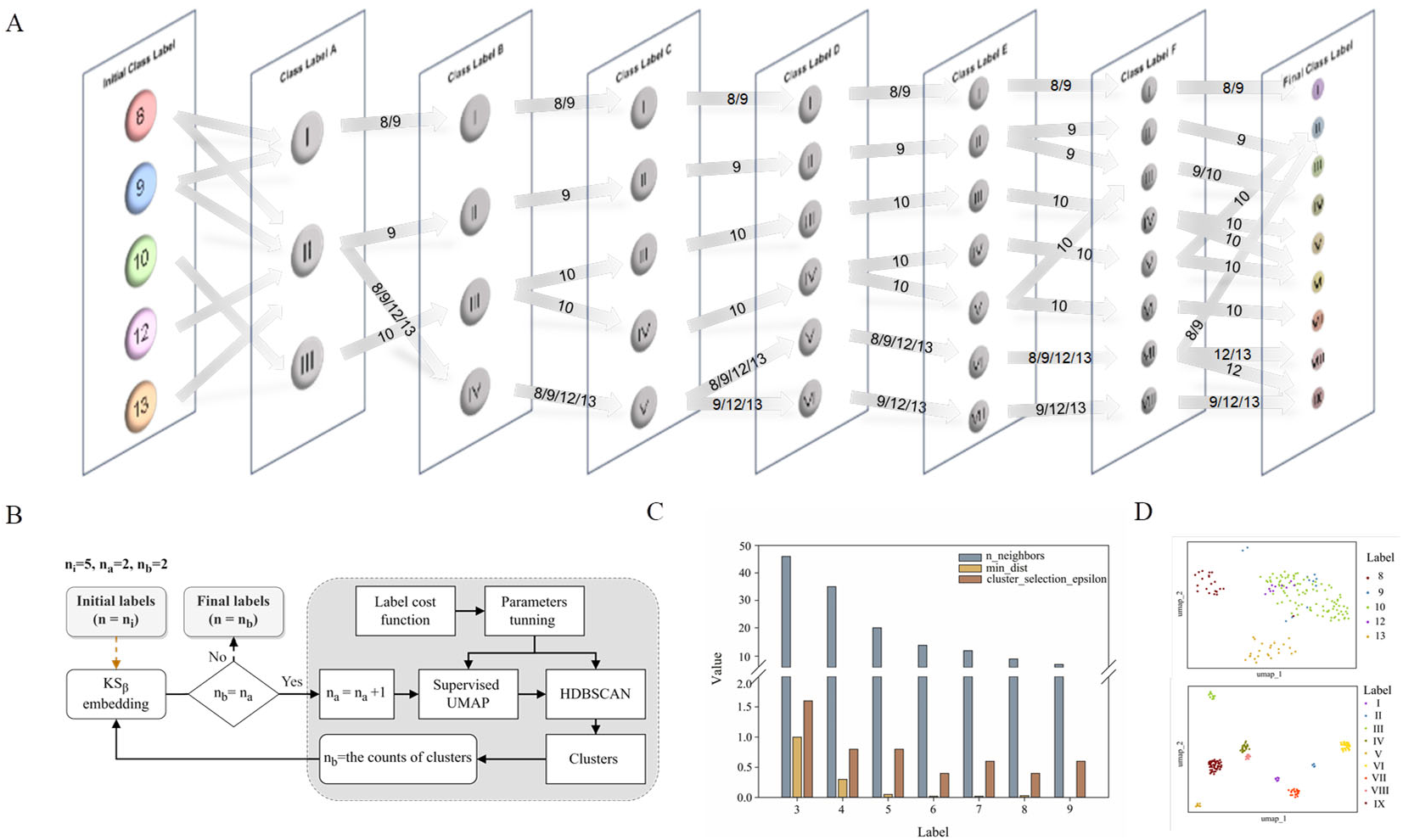
The process of T2PK class labeling. (**A**) Flowchart illustrating the process of KS_β_ labeling. Initial labels (n = 5) were generated based on the total number of T2PK building blocks, which included 8, 9, 10, 12 and 13 (refer to Fig. 1A). The algorithmic pipeline (Fig. 2B) was used to correct the inaccuracy of this classification. Certain class labels were bifurcated and merged, and ultimately, 9 new class labels were generated. (**B**) The process of KS_β_ labeling using supervised UMAP and HDBSCAN algorithms. Supervised UMAP and HDBSCAN were used as regularization techniques to constrain the latent distribution of KS_β_ embeddings and automatically tune UMAP and HDBSCAN hyperparameters using the labeling cost function to assign more appropriate class labels to embeddings. n_i_, the initial counts of labels; n_a_, the setting counts of labels; n_b_, the generated counts of labels (**C**) Parameter optimization of UMAP and HDBSCAN during the process of **A** and **B**. *n_neighbours* and *min_dist* parameters were automatically adjusted to refine the space and improve its alignment with the previous labels. Upon increasing the number of pre-defined labels, the approach tuned the *n_neighbours* and *min_dist* parameters to decrease their values, which allowed HDBSCAN to recognize and assign new labels to smaller and more localized features in the data space. (**D**) Supervised UMAP comparison of T2PK class labeling generated by five manually annotated class labels (up) and nine refined class labels (bottom).

As illustrated in Fig. 2A, the restructuring process of KS_β_ embeddings initially reset the counts of predefined labels to 3, followed by the automatic adjustment of the *n_neighbors* and *min_dist* parameters to refine the space and improve its alignment with these 3 class labels. HDBSCAN was then applied to cluster similar data points and assign new class labels to them. Certain previously assigned KS_β_ sequences were bifurcated and merged into a new label. For example, in the class label A column, label I is from partial initial label 8 and 9; label II is from partial initial label 8 and 9, and entire initial 12 and 13; and label III is from entire initial label 10 (Fig. 2A, Table S4). Upon increasing the number of predefined labels, the approach tuned the *n_neighbors* and *min_dist* parameters to decrease their values, which allowed HDBSCAN to recognize and assign new labels to smaller and more localized features in the data space (Fig. 2C, Fig.S3). This iterative process continued until the number of class labels assigned to the 164 KS_β_ embeddings reached 9, after which no more labels could be generated or some data points were identified as noise. To evaluate the distribution patterns of the 9 autogenerated labels, the supervised UMAP technique was employed again, revealing that these labels could represent the T2PK biosynthetic logics more accurately in the real world (Fig. 2D).

### Development of the T2PK classifier

As described in the previous session, four machine learning algorithms (random forest, XGBoost, SVM, and MLP with grid search hyperparameter tuning) were employed to train the initial T2PK classifiers on KS_β_ embeddings. The classifiers were trained on two distinct sets of class labels: one comprising five manually annotated class labels and another consisting of nine refined class labels (Fig. 2D). Due to the imbalanced nature of our dataset, the F1 score and confusion matrix were employed to assess and compare the performance of the classifiers trained on these two different sets of class labels. As outlined in Table 1, the MLP classifier yielded an F1 score of 0.88 on the test set when utilizing the nine refined class labels, whereas the F1 scores of the random forest, XGBoost, and SVM classifiers were 0.60, 0.82, and 0.92, respectively. Notably, the classifiers trained on the five manually annotated class labels exhibited inferior performance compared with those trained on the nine refined class labels.

**Table 1.** Performance metrics of initial and enhanced T2PK classifier with two types of class label trained by Random Forest, XGBoost, SVM, MLP. TPR: true positive rate; FPR: false positive rate.

| Classifier | Class type | TPR% | FPR% | Accuracy | Precision | Recall | F1-score |
| --- | --- | --- | --- | --- | --- | --- | --- |
| **Initial Classifier** |  |  |  |  |  |  |  |
| Random Forest | 5 classes | 46.00 | 9.12 | 0.79 | 0.49 | 0.46 | 0.46 |
|  | 9 classes | 59.72 | 3.40 | 0.76 | 0.66 | 0.60 | 0.60 |
| XGBoost | 5 classes | 56.00 | 8.41 | 0.82 | 0.72 | 0.56 | 0.60 |
|  | 9 classes | 83.33 | 2.01 | 0.85 | 0.82 | 0.83 | 0.82 |
| SVM | 5 classes | 74.00 | 4.48 | 0.88 | 0.90 | 0.74 | 0.78 |
|  | 9 classes | 96.76 | 0.71 | 0.94 | 0.91 | 0.97 | 0.92 |
| MLP | 5 classes | 71.00 | 4.44 | 0.88 | 0.84 | 0.71 | 0.76 |
|  | 9 classes | 92.59 | 1.60 | 0.88 | 0.87 | 0.93 | 0.88 |
| **Enhanced Classifier** |  |  |  |  |  |  |  |
| MLP | 5 classes | 80.00 | 2.25 | 0.94 | 0.96 | 0.80 | 0.84 |
|  | 9 classes | 94.44 | 0.89 | 0.94 | 0.98 | 0.94 | 0.95 |

To leverage the 2597 unlabeled KS_β_ embeddings, a consistency regularization-based semisupervised learning framework was adopted to train an enhanced MLP classifier based on the initial MLP classifier with two distinct sets of class labels. The enhanced MLP classifier was trained on both labeled and unlabeled data. In this process, the cross-entropy loss function was applied to the disturbed labeled data via Gaussian noise, while the mean-square error loss function was applied to both disturbed labeled and unlabeled data (Fig. 3A). This approach promoted the model to produce consistent predictions over time and resulted in a smoothed decision boundary, thereby enhancing the model’s generalization performance on unseen data and alleviating overfitting. A clear disturbance in the initial classifier at the beginning is shown in Fig. 3B. Overall, the performance evaluation demonstrated that the enhanced MLP classifier trained with nine refined class labels attained an F1 score of 0.95 on the test set, an increase of 0.07 compared with the initial one, indicating state-of-the-art performance.

**Fig. 3.**
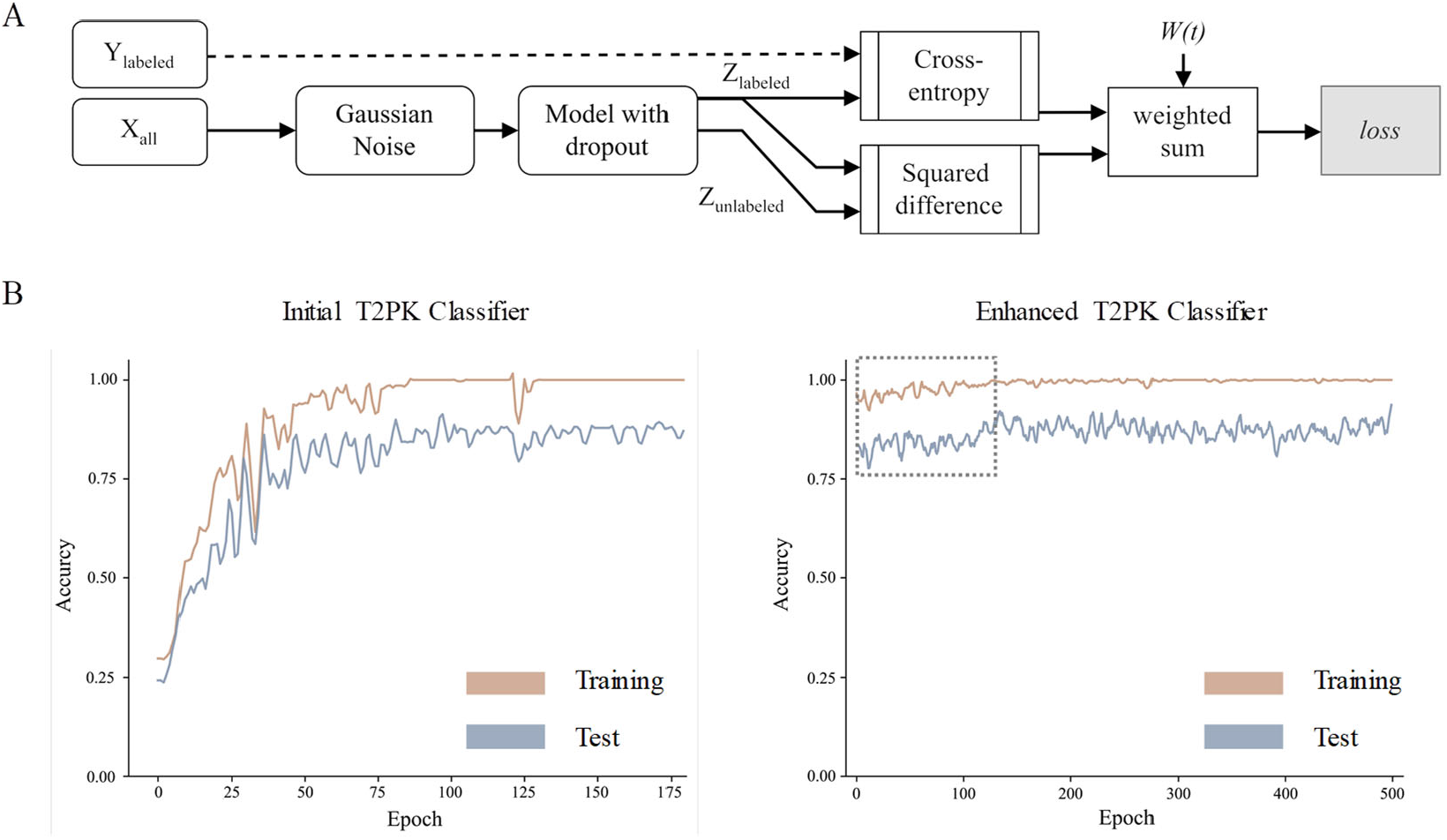
The development of T2PK classifier. (**A**) Flowchart depicting the consistency regularization-based semisupervised learning approach. Mean square errors were calculated for both labelled (164) and unlabeled (2597) data by adding Gaussian noise and applying model dropout with 50% feature loss. The cross-entropy loss was obtained using the initial and disturbed labeled data (164), while the weighted consistency regularization-based loss was obtained using the disturbed and undisturbed data, which included both labeled and unlabeled data (164 + 2597 = 2761). The total loss was calculated as the sum of cross-entropy loss and weighted consistency regularization-based loss. Note that the weight is proportional to training time. (**B**) Plots showing the accuracies of prediction for the initial (left) and enhanced (right) T2PK classifiers. Each epoch represents the relearning process of the entire shuffled dataset. The grey box with dash line indicates the changes in training and test accuracy in the initial phase after noise addition. Note that the enhanced T2PK classifier is developed based on the initial T2PK classifier.

### Out-of-distribution detection (ODD) for new T2PKs

Softmax-based classifiers have been criticized for generating overconfident posterior distributions when presented with ODD data (*27*). In this study, ODD data refer to KS_β_ sequences that do not belong to any of the nine refined classes. To overcome this limitation, we incorporated Mahalanobis distance-based scores (MDS) and anomaly detection techniques inspired by the generalized ODD framework (*28*). Specifically, we designated 164 KS_α_ sequences as ODD data and 164 KS_β_ sequences as in-distribution (ID) data and extracted feature vectors from the neural network consisting of input, hidden (n = 3), and output layers of the MLP classifier to compute the MDS for both ID and ODD data (Fig. 4A). We evaluated the classification performance of each layer using a one-class SVM on the ID and ODD datasets and found that the hidden layer 1 exhibited superior performance in detecting ODD data (Fig. 4B, Fig. S4).

**Fig. 4.**
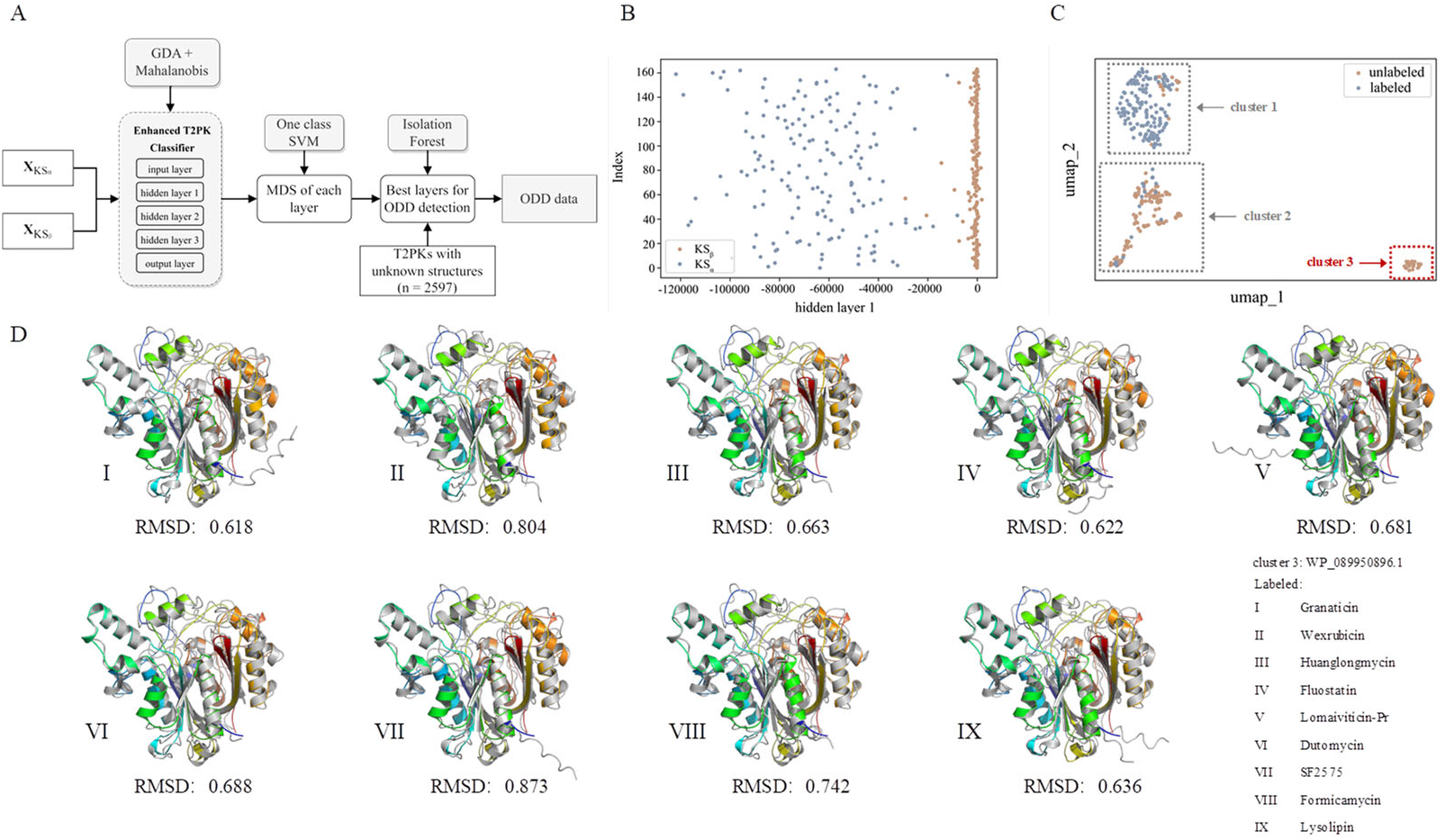
Detection of novel T2PKs using ODD framework. (**A**) Workflow showing the process for detecting novel T2PKs; KS_α_ and KS_β_ as ODD and ID data respectively derived from a single biosynthetic gene cluster were both used as input to MLP classifier. MDS system was employed to measure the distance between individual data points of which the feature vectors were extracted from the input, hidden and output layers of the MLP classifier. The classification performance of each layer was evaluated by one-class SVM. An isolation forest model was trained using the generated MDS data from the best layer and furthermore the larger dataset of 2597 unlabeled data points were analysed to detect abnormal data which is defined as “novel” in this work. (**B**) The classification performance of the hidden layer 1. For the performance of the other layers, see Fig. S4. (**C**) Cluster 3 containing the KS_β_ that most likely involve the biosynthesis of novel T2PKs. (**D**) Representative in-silico predictions of the protein structures generated by ESMFold. Grey structure indicates the labelled KS_β_ for the biosynthesis of known T2PK, while colour structures indicate unlabeled KS_β_ from cluster 3 in **C**, respectively.

Next, we used the generated MDS data from hidden layer 1 to train an isolation forest model and subsequent detect abnormal data within a larger dataset of 2597 unlabeled data points. Setting the contamination parameter of isolation forest to 0.01, combining the labeled and abnormal datasets, and performing UMAP dimension reduction, we identified three ODD clusters consisting of overall 153 abnormal data. On the one hand, certain data points from ODD cluster 1 and 2 were grouped within the labeled dataset (grey dotted box in Figure 4C). This does not mean the inaccuracy of the detection, rather it indicates the KS_β_ from these two clusters still have some similar embeddings with the 164 labeled KS_β_ and suggests their corresponding chemical structures may possess some common characters with those structurally known T2PKs. On the other hand, we observed that ODD cluster 3 consisting of 20 sequences was entirely separated from the labeled dataset and located at a considerable distance (Figure 4C). Our hypothesis is that the KS_β_ proteins in this ODD cluster may present novel protein structures which differ from the previously labeled KS_β_ proteins, and that these novel structures probably involve in the biosynthesis of novel T2PKs. To test this hypothesis, we conducted in-silico predictions of the protein structures for all data points within this cluster using ESMFold and calculated the root mean square deviation (RMSD) between the predicted structures of the 20 unlabeled and 164 labeled KS_β_ proteins. The protein structures belonging to the ODD cluster 3 and those within IV, V, VI, and VII classes were found to exhibit at an average RMSD value of 0.59 Å (Figure S5 and Table S5). Table S6 shows the calculated RMSD values between ODD cluster 3 and IV-VII classes, which allows to preset a threshold value at 0.4 to distinguish intra-class and inter-class. This means for novel T2PKs exploration, attentions should be focused on KS_β_ proteins with RMSD value between the ODD cluster and other classes over 0.4, as illustrated in Figure 4D.

### Predicting T2PKs from bacterial genomes

Thus far, we have shown that DeepT2 is capable of effectively accelerating the identification of T2PKs. We therefore wanted to investigate the advantages of this model to reveal the potential of any T2PKs produced by the actinomycetes isolated or stocked in our laboratory. As such, we sequenced 6 *Streptomyces* strains and confirmed their genomic independence through average nucleotide identity comparison. Subsequently, we utilized DeepT2, DeepBGC, and antiSMASH (*29*) to predict the T2PK produced from these 6 strains. Our results indicate that DeepT2 outperforms DeepBGC and antiSMASH in terms of predicting T2PKs, as the latter two tools only identify the BGCs. Furthermore, we examined the capacity of these tools to handle metagenomic sequences by mixing the 6 genomes and submitting them to above tools. The result showed that only DeepT2 can handle metagenomic input, with the output results being identical to those obtained from single-genome input. However, neither web server nor local antiSMASH allows for direct submission of metagenomic sequences. Additionally, we evaluated the performance of these tools on single-gene input by extracting candidate KS_β_ sequences from the 6 genomes and submitting them as a single sequence to the above tools, and found only DeepT2 supported single-gene input and made accurate predictions (Table S7).

As described before, DeepT2 employs a multi-step approach that involves first predicting T2PK class labels based on input KS_β_ sequences, then calculating the Euclidean distance between ID samples in a 3-dimensional space, and finally using the distance to determine the similarity between the input data and the ID sample (Fig. 5). In this way, we selected three top closest ID sample as predicted T2PK for the unknown KS_β_ input. As an overall result, 12 KS_β_ protein sequences were detected from 6 *Streptomyces* genome that fell into four classes (Table S7), and the corresponding T2PKs were closest to alnumycin/granaticin/frenolicin in class I (*30-32*), polyketomycin/dutomycin/arimetamycin (*33-35*) in class VI, fasamycin/formicamycin/Sch in class VIII (*36-38*), and lysolipin/rubromycin/anthrabenzoxocinone in class IX (*39-41*), respectively (Fig. 5, Table S7). To confirm this prediction, the strains were then inoculated and their metabolites were analyzed by liquid chromatography high-resolution mass spectrometry. As the result, alnumycin from WY86, polyketomycin from WY13 and lysolipin from PS14 were observed (Fig. 5, Fig. S6), whereas fasamycin from *S. kanamyciticus* was not detected under laboratory conditions as described in a recent study (*42*).

**Fig. 5.**
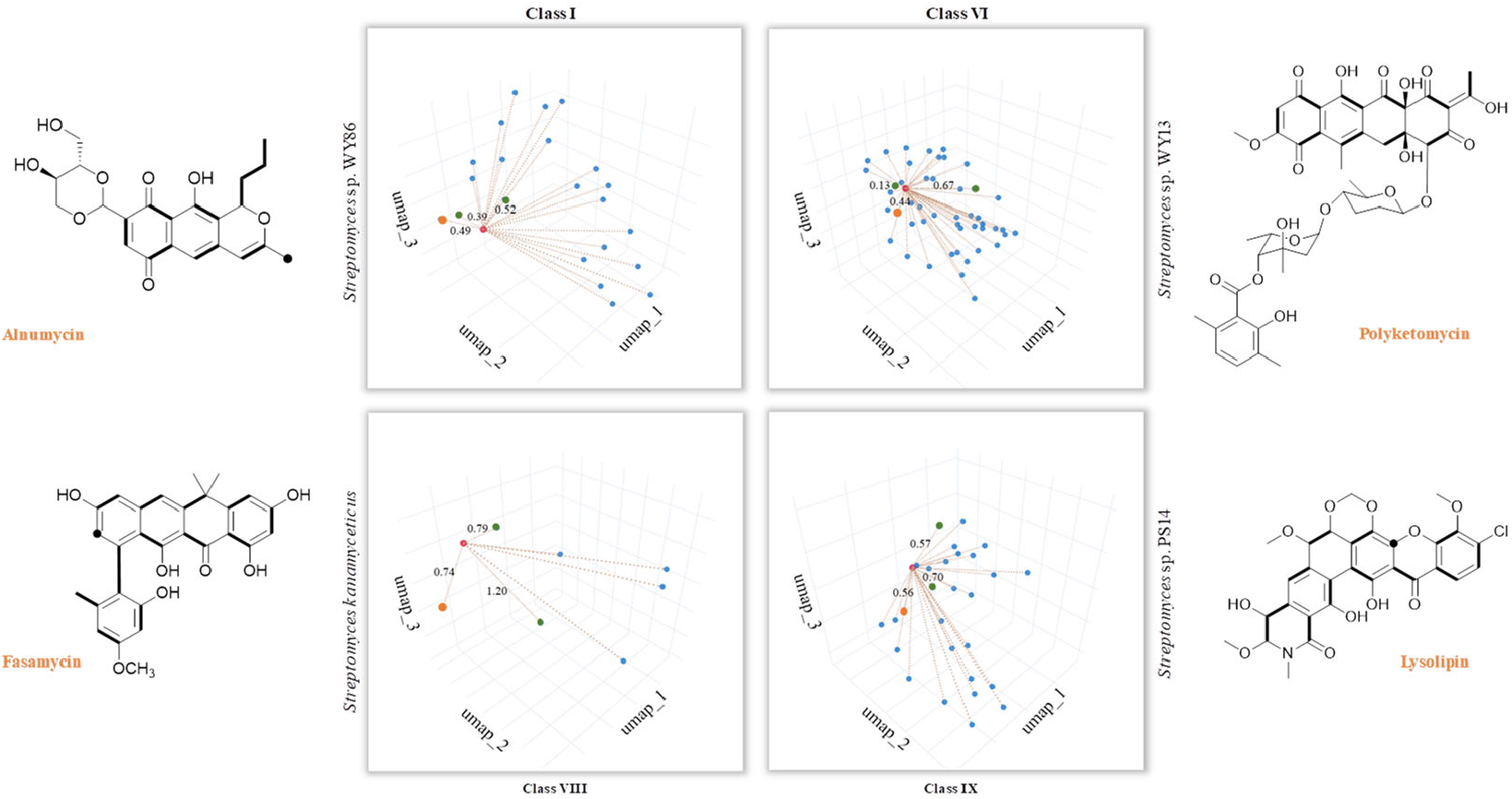
T2PK prediction from bacterial genomes. DeepT2 was performed and the identified compounds alnumycin A, polyketomycin, and lysolipin were subsequently confirmed via high-resolution mass spectra (Fig. S6). Euclidean distances for each predicted candidate with top 3 similar T2PKs were annotated beside the dash lines (red, T2PKs to be detected; orange, the most similar T2PKs experimentally confirmed; green, other similar T2PKs that have close KS_β_ structures but different natural products). Further information can be found in Table S7. The ground-truth T2PK of bacteria is denoted by the orange point in each figure. Bold bonds in each chemical structure indicate the building block units incorporated into the polyketide backbone, while the black dot indicates a single carbon from the build block unit in which the adjacent carbon from the same build block is lost during the polyketide biosynthesis via decarboxylation.

## Discussion

T2 polyketide synthase is a family of single heterodimeric ketosynthases that iteratively catalyzes the elongation of the polyketide chain structure, leading to our inability to precisely predict T2PK structures. As introduced previously, despite the multiple sequence alignment approaches based on KS_β_ (*5, 6*), incorporation of new sequences into the evolutionary model may alter the structure of the original phylogenetic tree and therefore compromise the accuracy of the predictions. To address this issue, we propose DeepT2, an end-to-end deep learning strategy, to directly identify T2PKs from bacterial genomes. Leveraging the concept of natural language processing, our approach embeds KS_β_ as feature vectors, enabling the representation of protein structural information. We employ semisupervised learning to link KS_β_ embedding vectors with compound labels, facilitating the rapid identification of known and novel T2PKs. Importantly, in contrast to other BGC prediction tools, such as DeepBGC (*17*) and antiSMASH, which require complete T2PK BGC sequences, our DeepT2 model can accurately predict T2PK categories using only KS_β_ sequences. Furthermore, our novelty detection framework embedded in DeepT2 has promising potential for identifying new KS_β_. However, a question that may arise is that considering the limited biological data volume, is it still possible to use it to develop advanced algorithms even without big data? This is because in the biological research field, classic machine learning and currently popular deep learning are both hampered by poor accuracy or overfitting caused by the smaller training dataset. Indeed, during model development, we had debated whether the proof of concept should be data-centered or training-centered. Fortunately, the ensemble method using a small data volume based on pretraining and semisupervised learning seems to be a promising solution, at least for this work. In addition, as with other machine learning algorithms, DeepT2 is expected to improve as more KS_β_ sequences are discovered in microbial genomes over time.

The task of few-shot supervised learning requires an approach that transcends traditional supervised neural networks (*43*). In this context, our work adopts the concept of transfer learning, where ESM-2 is utilized to explore the connection between KS_β_ embeddings and T2PK structures. While the dimension reduction results indeed indicate that the embedding vectors obtained by ESM-2 closely fit with the compound class labels, it is important to note that certain embeddings still necessitate further labeling refinement and correction. Consequently, we performed label reconstruction using supervised UMAP instead of unsupervised UMAP to ensure that the resulting reduced-dimensional space consistently captures the features of the compound skeleton. This approach differs from traditional unsupervised learning for clustering (*44*), as it strives to strike a balance between the sequence embeddings and the compound class labels to improve the model’s accuracy. For example, T2PK AQ-256-8 consists of 8 building blocks, but its KS_β_ is confirmed as ancestral nonoxidative, which differs from other KS_β_s that involve the biosynthesis of T2PKs with 8 building blocks (*6*). Clearly, the state-of-the-art performance of the model trained with 9 refined class labels suggests that the classification effect is unsatisfactory when simply using five biosynthetic building blocks as labels. This finding suggests that KS_β_ not only affects the counts of building blocks but also determines a rough topology prior to cyclization or aromatization. To the best of our knowledge, this is the first algorithm for T2PK classification and prediction in such a manner, which, as an alternative to sophisticated protein sequence alignment, might showcase a paradigm shift in genome-mining approaches for natural product discovery.

As shown above, we improved the generalization ability of the softmax-based T2PK classifier by employing a consistency-regularization-based semisupervised learning framework that utilized 2597 KS_β_ whose corresponding natural product structures currently remain unknown. However, such models may demonstrate overconfidence in discerning novel KS_β_ sequences in the open world (*27*). To address this concern, an ODD framework based on the Mahalanobis distance was implemented for multiclass novelty detection (*45*). Notably, certain samples (from 2597 KS_β_ sequences) are proximal to the labeled data (from 164 KS_β_ sequences) because such labeled T2PKs with entirely novel carbon skeletons have only been discovered in recent years, such as formicamycin (*36*) and dendrubin (*46*). Therefore, to avoid false positives in novelty detection, we selected only 20 potential new class samples that are distant from the labeled samples for demonstration. Greater details regarding the enzymatic information and chemical structures for these T2PKs will be studied in future work.

This study demonstrates the capacity of DeepT2 to predict T2PKs from single or mixed genomic datasets. However, some limitations must be acknowledged. While the training data included bacterial genomes from different phyla, certain biases may hinder the model’s ability to detect novel T2PKs in poorly characterized bacterial sources within complex microbiomes. Although the model was validated with *Streptomyces* genomes as a showcase in this study, expanding the bacterial genome resources is crucial to improve the model’s overall performance. Additionally, the current version of DeepT2 is capable of predicting T2PKs from single genes as input, but it requires complete sequences of at least 300 amino acids (the average length of KS_β_ is around 400 amino acid). For predicting other tailoring modifications, such as methylation or halogenation, supplementing DeepT2 with antiSMASH or DeepBGC is recommended. Nonetheless, despite these limitations, the DeepT2 model outperforms other methods and represents a valuable algorithm for KS_β_ identification and T2PK discovery. This study also inspires future research to identify which catalytic domains in KS_β_ contribute to chemical differences through PLM and thus provides more insights into the KS_β_ evolution and T2KS biosynthetic mechanisms, and this is currently ongoing in our laboratory. Moreover, as the application of language models in prompt tuning for zero-shot prediction, as well as the generative models such as autoregressive neural networks is gradually emerging (*47-49*), we are now prompted to explore such models for KS_β_ studies. We therefore anticipate that this work will aid in the application of genome mining approaches to discover new KS_β_ and novel T2PKs and have important clinical implications for transforming microbiome data into therapeutic interventions.

## Materials and methods

### Protein sequence data preparation and embedding using protein language models

A dataset comprising 164 KS_β_ sequences with known chemical structures was retrieved from the NCBI database. These sequences were designated as ‘labeled’. In contrast, KS_β_ sequences without corresponding chemical structures were classified as ‘unlabeled’. To further curate the unlabeled KS_β_, the 164 labeled KS_α_ and KS_β_ sequences were used to obtain 20 kb putative T2PK minimum BGCs following the in-house pipeline described by Chen *et al. (6*). For the criterion of a reliable T2PK gene cluster, KS_α_ and KS_β_ sequences should be identified in the same contig. This process yielded 2597 KS_β_ sequences without corresponding chemical structures. Additionally, the non-KS_β_ sequences from actinobacteria were obtained from UniRef50, with a focus on sequences between 300 and 500 amino acids in length. To eliminate irrelevant entries, we excluded any sequence matching the keywords ‘beta ketoacyl synthase’ or ‘chain length factor’. Taken together, our dataset consisted of 2761 (164 + 2597) KS_β_ sequences and 761,334 non-KS_β_ sequences, which were subjected to subsequent analysis.

As protein sequences should be represented with numerical vectors before being input to the classification algorithm, we employed five general protein language models (PLMs) to embed the amino acids of each sequence, resulting in a representation vector obtained by averaging the embeddings across the entire sequence. We also aimed to investigate which PLMs produced high-quality learned representations for distinguishing functions, as well as whether these representations were associated with the given labels. To achieve this, we used the UMAP dimension-reduction algorithm to visualize the high-dimensional embeddings. Our results demonstrated that the ESM-2 model, with 3 billion parameters, outperformed the other models in terms of learned representation quality. This finding suggests that ESM-2 may be particularly useful for protein sequence classification tasks.

### Binary KS_β_ classifier development

To construct a robust binary classification model, we utilized an 80/10/10 split of the KS_β_ and non-KS_β_ datasets to produce a training, validation and test set, respectively. We trained four classifiers, namely, random forest, XGBoost, support vector machine (SVM) using the scikit-learn Python module (version 1.2) and multilayer perceptron (MLP) using the PyTorch module (version 2.0). The selected hyperparameters for each classifier are documented in Table S8. For the MLP classifier, a dropout layer with a value of 0.5 was incorporated into the network, and a binary cross-entropy loss function was utilized. To evaluate the performance of the models, a 5-fold cross-validation technique was implemented, and the accuracy value of each fold was averaged. Due to the imbalanced nature of the two datasets, only accuracy, confusion matrix, and F1 score were employed as evaluation metrics for the final model performance. The scikit-learn Python module was used to compute all these metrics.

### Relabeling 164 KS_β_ protein sequences

Initially, 164 KS_β_ sequences were categorized into 5 classes based on the number of building blocks for the main carbon skeleton. Next, we developed a pipeline for tuning hyperparameters in clustering using HDBSCAN (*50*) and UMAP (*51*) to refine and expand the predefined 5 classes labels. Our pipeline was guided by the chatintents python packages (https://github.com/dborrelli/chat-intents), but we replaced unsupervised UMAP with supervised UMAP. This pipeline comprised four major functions: *generate_clusters, score_clusters, objective* and *bayesian_search*. Briefly, the main function *bayesian_search* takes in a dataset, hyperparameter space and other parameters. It deploys the ‘trials’ object within the *bayesian_search* function to monitor the results of each evaluation of the *objective* function. The *objective* function, on the other hand, receives hyperparameters from the space, applies the *generate_clusters* function to create a clustering object, and computes the number of clusters and a cost metric via the *score_clusters* function. It also imposes a penalty on the cost if the number of clusters falls outside the desired range. The Bayesian optimization process is conducted using the *fmin* function from the *hyperopt* package, with the *tpe*.*suggest* algorithm eventually yielding the optimal hyperparameters found, the resulting clustering object, and the trials object for further scrutiny. The hyperparameter space encompasses diverse parameters, including *n_neighbors* and *min_dist* of UMAP and *cluster_selection_epsilon* of HDBSCAN. The detailed range of hyperparameter spaces is supplemented in Table S9. Overall, the pipeline was designed to optimize the creation of new local clusters from global clusters, utilizing supervised UMAP and HDBSCAN with the aid of label penalty and Bayesian optimization techniques. By following this pipeline, we are able to generate new class labels based on the previous class labels while simultaneously improving their consistency with the protein structure information that is embedded in the data.

### T2PK classifier architecture and training in different label types

In the present investigation, a total of 164 KS_β_ sequences, each with known chemical structures, were assigned five manually annotated and nine autogenerated class labels. Owing to the significant imbalance in the labeled dataset, the dataset was only divided into training and test sets following an 80/20 ratio. Notably, random partitioning of data is typically avoided in biological sequence modeling due to its tendency to yield an overly simplistic evaluation of generalization. Thus, sticking to the 80/20 ratio, we conducted manual partitioning of the dataset. Detailed information is provided in Table S4. To assess the quality of the assigned labels and to determine the most suitable multiclass classification algorithm, four multiclass classifiers were trained using the scikit-learn and Pytorch modules. Hyperparameter optimization was performed using grid searches, and model performance was assessed by the metrics mentioned above.

To enhance the robustness of the initial MLP classifier, we employed a consistency regularization-based semisupervised framework, as proposed by Laine et al (*52*). The loss function comprised two distinct components: the standard cross-entropy loss, which exclusively evaluated the labeled inputs, and a second component that evaluated all inputs (both labeled and unlabeled). This second component penalized divergent predictions for the same training input, measured by the mean square difference between the prediction vectors and perturbed vectors. To augment the unlabeled data, Gaussian noise was added to the embeddings by specifying a standard deviation for the Gaussian distribution. The optimal amount of noise was determined based on its effect on the training accuracy. The resulting neural network classifier was then trained using a weighted total loss function, consisting of both the consistency loss and the standard cross-entropy loss. The detailed hyperparameter spaces mentioned above have been summarized in Table S8.

### Detection of T2PKs with potentially new skeletons

Softmax-based classifiers are known to generate overconfident posterior distributions when presented with out-of-distribution detection (ODD) data. To address this issue, a generative classifier, specifically Gaussian discriminant analysis, and a Mahalanobis distance-based framework were utilized to obtain confidence scores, as described by Lee et al (*45*). To evaluate the performance of this approach, 164 KS_α_ sequences were designated ODD data, and 164 KS_β_ sequences were designated in-distribution (ID) data. The feature vector for each layer was extracted from the neural network, and a generative classifier was applied to obtain the mean and covariance matrix for each class. The Mahalanobis distance-based scores were then calculated between a test sample and the closest class Gaussian to obtain confidence scores for both datasets. To identify the feature layer that was most suitable for distinguishing between ID and ODD data, all layers were evaluated using a one-class SVM on both datasets. The layer that exhibited the best performance on both datasets was selected. The isolation forest, a general abnormal data detection algorithm, was then used to detect novelty data from an unlabeled dataset. The protein structure of the detected KS_β_ novelty data was predicted using ESMFold (*20*) *in silico*, and the difference in protein structures was determined using the root-mean-square deviation.

### General microbiology and chemistry experiments

Total DNA of six *Streptomyces* strains was used for experimental confirmation, and their genomes were extracted using a custom genomic extraction protocol. The genomes were sequenced using a one-dimensional MinION flow cell with an r9.4.1 flow cell from Oxford Nanopore Technologies, UK, and base-calling was performed with Guppy v6.2.1. Additionally, some DNA samples were sequenced using MiSeq sequencing from Illumina, CA, USA. Quality control of the long reads (LRs) and short reads (SRs) was conducted using fastqc v0.11.6 (*53*), with quality control performed using different software suites: Porechop v0.2.4 (*54*) and Filtlong v0.2.0 (https://github.com/rrwick/Filtlong) for LRs and FASTX-Toolkit for SRs. A hybrid assembly using Unicycler v0.5.0 (*55*) was then performed by combining LRs and SRs. For samples with LRs only, the assembly was carried out using canu 2.1 (*56*) and polished with racon 1.0 (*57*). The protein sequences of all genomes were predicted using prokka v1.14.6 (*58*). The BGCs of the six genomes were predicted using DeepBGC (*17*) and antiSMASH (*29*). The performance of DeepT2, DeepBGC and antiSMASH in predicting T2PKs was compared.

Unless stated otherwise, all chemicals were supplied by Macklin. All solvents were of HPLC grade or equivalent. Actinomycetes were cultivated at 30 °C on ISP2 agar. Crude metabolites were extracted by ethyl acetate. Samples were analyzed by LCMS/MS on an Agilent G6500 UHPLC system attached to a quadrupole time-of-flight (Q-ToF) mass spectrometer. The spray chamber conditions were as follows: nebulizer, 5 L/min; drying gas, 200; sheath gas temperature, 350 °C; sheath gas flow, 11 L/min; and drying gas on, 5 L/min. The instrument was calibrated using an API-TOF Reference Mass Solution Kit according to the manufacturer’s instructions. The following analytical LCMS method was used throughout this study: Phenomenex Kinetex C18 column (100 × 2.1 mm, 100 Å); mobile phase A: water + 0.1% formic acid; mobile phase B: acetonitrile + 0.1% formic acid. Elution gradient: 0-1 min, 20% B; 1-12 min, 20%-100% B; 12-14 min, 100% B; 14-14.1 min, 100%-20% B; 14.1-17 min, 20% B; flow rate: 0.3 mL/min; injection volume: 10 µL. HPLC was performed on an Agilent 1290 system, and the following method was used throughout this study: Phenomenex Kinetex C18 column (100 × 2.1 mm, 100 Å); mobile phase A: water + 0.1% formic acid; mobile phase B: acetonitrile + 0.1% formic acid. Elution gradient: 0-1 min, 20% B; 1-12 min, 20%-100% B; 12-14 min, 100% B; 14-14.1 min, 100%-20% B; 14.1-17 min, 20% B; flow rate: 0.3 mL/min; injection volume: 10 µL. UV: 250 and 415 nm.

### Molecular networking description

A molecular network was created using the online workflow on the GNPS website (http://gnps.ucsd.edu) (*59*). The data were filtered by removing all MS/MS fragment ions within +/- 17 Da of the precursor m/z. MS/MS spectra were window filtered by choosing only the top 6 fragment ions in the +/- 50 Da window throughout the spectrum. The precursor ion mass tolerance was set to 1 Da with an MS/MS fragment ion tolerance of 0.5 Da. A network was then created where edges were filtered to have a cosine score above 0.2 and more than 2 matched peaks. Furthermore, edges between two nodes were kept in the network if and only if each of the nodes appeared in each other’s respective top 10 most similar nodes. Finally, the maximum size of a molecular family was set to 100, and the lowest scoring edges were removed from molecular families until the molecular family size was below this threshold.

## Data availability

The authors declare that the data, materials and code supporting the findings reported in this study are available from the authors upon reasonable request. The DeepT2 is available at GitHub repository https://github.com/Qinlab502/deept2.

## Authors contributions

Zhiwei Qin and Heqian Zhang designed and supervised the research, Jiaquan Huang and Qiandi Gao performed the bioinformatic and computing analysis, Yaxin Wu performed the microbial fermentation and chemical analysis, Ying Tang participated the algorithm development. All authors analysed and discussed the data. Zhiwei Qin, Heqian Zhang and Jiaquan Huang wrote the manuscript and all authors commented.

## Acknowledgements

This work was supported by the National Natural Science Foundation of China (32170079, 32200035 and 12105014), the Natural Science Foundation of Guangdong (2021A1515012026), Guangdong Talent Scheme (2021QN020100), Beijing Normal University via the Youth Talent Strategic Program Project (310432104), as well as Guangdong Innovation Research Team for Plant-Microbe Interaction. We also thank the Interdisciplinary Intelligence Super Computer Center, Beijing Normal University at Zhuhai, for High Performance Computing for access to computational resources.

## Reference

1. C. Hertweck, A. Luzhetskyy, Yu. Rebets and A. Bechthold. Nat. Prod. Rep 24, 162 (2007).

2. S.-C. Tsai, The structural enzymology of iterative aromatic polyketide synthases: a critical comparison with fatty acid synthases. Annual Review of Biochemistry 87, 503–531 (2018).

3. C. Hertweck, The biosynthetic logic of polyketide diversity. Angewandte Chemie International Edition 48, 4688–4716 (2009).

4. A. Bräuer, Q. Zhou, G. L. Grammbitter, M. Schmalhofer, M. Rühl, V. R. Kaila, H. B. Bode, M. Groll, Structural snapshots of the minimal PKS system responsible for octaketide biosynthesis. Nature Chemistry 12, 755–763 (2020).

5. M. E. Hillenmeyer, G. A. Vandova, E. E. Berlew, L. K. Charkoudian, Evolution of chemical diversity by coordinated gene swaps in type II polyketide gene clusters. Proceedings of the National Academy of Sciences 112, 13952–13957 (2015).

6. S. Chen, C. Zhang, L. Zhang, Investigation of the molecular landscape of bacterial aromatic polyketides by global analysis of type II polyketide synthases. Angewandte Chemie International Edition 61, e202202286 (2022).

7. C. P. Ridley, H. Y. Lee, C. Khosla, Evolution of polyketide synthases in bacteria. Proceedings of the National Academy of Sciences 105, 4595–4600 (2008).

8. J. Kim, G.-S. Yi, PKMiner: a database for exploring type II polyketide synthases. BMC microbiology 12, 1–12 (2012).

9. R. Villebro, S. Shaw, K. Blin, T. Weber, Sequence-based classification of type II polyketide synthase biosynthetic gene clusters for antiSMASH. Journal of Industrial Microbiology Biotechnology 46, 469–475 (2019).

10. E. C. Alley, G. Khimulya, S. Biswas, M. AlQuraishi, G. M. Church, Unified rational protein engineering with sequence-based deep representation learning. Nature methods 16, 1315–1322 (2019).

11. A. Elnaggar, M. Heinzinger, C. Dallago, B. Rost, End-to-end multitask learning, from protein language to protein features without alignments. bioRxiv, 864405 (2019).

12. N. J. Merwin, W. K. Mousa, C. A. Dejong, M. A. Skinnider, M. J. Cannon, H. X. Li, K. Dial, M. Gunabalasingam, C. Johnston, N. A. Magarvey, DeepRiPP integrates multiomics data to automate discovery of novel ribosomally synthesized natural products. P Natl Acad Sci USA 117, 371–380 (2020).

13. C. Rios-Martinez, N. Bhattacharya, A. P. Amini, L. Crawford, K. K. Yang, Deep self-supervised learning for biosynthetic gene cluster detection and product classification. bioRxiv, 2022.2007. 2022.500861 (2022).

14. Y. Ma, Z. Guo, B. Xia, Y. Zhang, X. Liu, Y. Yu, N. Tang, X. Tong, M. Wang, X. Ye, Identification of antimicrobial peptides from the human gut microbiome using deep learning. Nat Biotechnol 40, 921–931 (2022).

15. Y. Tang, A. Hoffmann, Quantifying information of intracellular signaling: progress with machine learning. Rep. Prog. Phys 85, 086602 (2022).

16. L. Yann, B. Yoshua, H. Geoffrey, Deep learning. Nature 521, 436–444 (2015).

17. G. D. Hannigan, D. Prihoda, A. Palicka, J. Soukup, O. Klempir, L. Rampula, J. Durcak, M. Wurst, J. Kotowski, D. Chang, R. R. Wang, G. Piizzi, G. Temesi, D. J. Hazuda, C. H. Woelk, D. A. Bitton, A deep learning genome-mining strategy for biosynthetic gene cluster prediction. Nucleic Acids Research 47, p(2019).

18. A. Rives, J. Meier, T. Sercu, S. Goyal, Z. Lin, J. Liu, D. Guo, M. Ott, C. L. Zitnick, J. Ma, Biological structure and function emerge from scaling unsupervised learning to 250 million protein sequences. Proceedings of the National Academy of Sciences 118, e2016239118 (2021).

19. S. Unsal, H. Atas, M. Albayrak, K. Turhan, A. C. Acar, T. Doğan, Learning functional properties of proteins with language models. Nature Machine Intelligence 4, 227–245 (2022).

20. Z. Lin, H. Akin, R. Rao, B. Hie, Z. Zhu, W. Lu, N. Smetanin, R. Verkuil, O. Kabeli, Y. Shmueli, Evolutionary-scale prediction of atomic-level protein structure with a language model. Science 379, 1123–1130 (2023).

21. F. Teufel, J. J. Almagro Armenteros, A. R. Johansen, M. H. Gíslason, S. I. Pihl, K. D. Tsirigos, O. Winther, S. Brunak, G. von Heijne, H. Nielsen, SignalP 6.0 predicts all five types of signal peptides using protein language models. Nat Biotechnol 40, 1023–1025 (2022).

22. M. H. Hoie, E. N. Kiehl, B. Petersen, M. Nielsen, O. Winther, H. Nielsen, J. Hallgren, P. Marcatili, NetSurfP-3.0: accurate and fast prediction of protein structural features by protein language models and deep learning. Nucleic Acids Research 50, W510–W515 (2022).

23. H. Song, M. Kim, D. Park, Y. Shin, J.-G. Lee, Learning from noisy labels with deep neural networks: A survey. IEEE Transactions on Neural Networks Learning Systems, (2022).

24. Y. Ouali, C. Hudelot, M. Tami, An overview of deep semi-supervised learning. arXiv preprint arXiv:.05278, (2020).

25. K. Lee, K. Lee, H. Lee, J. Shin, A simple unified framework for detecting out-of-distribution samples and adversarial attacks. arXiv:1807.03888, (2018).

26. L. Zhou, S. Pan, J. Wang, A. V. Vasilakos, Machine learning on big data: Opportunities and challenges. Neurocomputing 237, 350–361 (2017).

27. A. Nguyen, J. Yosinski, J. Clune, in Proceedings of the IEEE conference on computer vision and pattern recognition. (2015), pp. 427–436.

28. J. Yang, K. Zhou, Y. Li, Z. Liu, Generalized out-of-distribution detection: A survey. arXiv preprint arXiv:.11334, (2021).

29. B. Kai, S. Simon, K. Alexander M, C.-P. Zach, V. W. Gilles P, M. Marnix H, W. Tilmann, antiSMASH 6.0: improving cluster detection and comparison capabilities. Nucleic acids research 49, W29–W35 (2021).

30. T. Oja, L. Niiranen, T. Sandalova, K. D. Klika, J. Niemi, P. Mantsala, G. Schneider, M. Metsa-Ketela, Structural basis for C-ribosylation in the alnumycin A biosynthetic pathway. Proc Natl Acad Sci U S A 110, 1291–1296 (2013).

31. K. Ichinose, D. J. Bedford, D. Tornus, A. Bechthold, M. J. Bibb, W. P. Revill, H. G. Floss, D. A. Hopwood, The granaticin biosynthetic gene cluster of Streptomyces violaceoruber Tü22: sequence analysis and expression in a heterologous host. Chemistry biology 5, 647–659 (1998).

32. C. Han, Z. Yu, Y. Zhang, Z. Wang, J. Zhao, S.-X. Huang, Z. Ma, Z. Wen, C. Liu, W. Xiang, Discovery of frenolicin B as potential agrochemical fungicide for controlling fusarium head blight on wheat. Journal of Agricultural Food Chemistry 69, 2108–2117 (2021).

33. M. Daum, I. Peintner, A. Linnenbrink, A. Frerich, M. Weber, T. Paululat, A. Bechthold, Organisation of the biosynthetic gene cluster and tailoring enzymes in the biosynthesis of the tetracyclic quinone glycoside antibiotic polyketomycin. ChemBioChem 10, 1073–1083 (2009).

34. L.-J. Xuan, S.-H. Xu, H.-L. Zhang, Y.-M. Xu, M.-Q. Chen, Dutomycin, a new anthracycline antibiotic from Streptomyces. The Journal of antibiotics 45, 1974–1976 (1992).

35. H. S. Kang, S. F. Brady, Arimetamycin A: improving clinically relevant families of natural products through sequence-guided screening of soil metagenomes. Angewandte Chemie International Edition 52, 11063–11067 (2013).

36. Z. Qin, J. T. Munnoch, R. Devine, N. A. Holmes, R. F. Seipke, K. A. Wilkinson, B. Wilkinson, M. I. Hutchings, Formicamycins, antibacterial polyketides produced by Streptomyces formicae isolated from African Tetraponera plantants. Chemical science 8, 3218–3227 (2017).

37. Z. Qin, R. Devine, T. J. Booth, E. H. Farrar, M. N. Grayson, M. I. Hutchings, B. Wilkinson, Formicamycin biosynthesis involves a unique reductive ring contraction. Chemical Science 11, 8125–8131 (2020).

38. G. Blanco, P. Brianb, A. Pereda, C. Mendez, J. Salas, K. F. Chater, Hybridization and DNA sequence analyses suggest an early evolutionary divergence of related biosynthetic gene sets encoding polyketide antibiotics and spore pigments in Streptomyces spp. Gene 130, 107–116 (1993).

39. P. Lopez, A. Hornung, K. Welzel, C. Unsin, W. Wohlleben, T. Weber, S. Pelzer, Isolation of the lysolipin gene cluster of Streptomyces tendae Tü 4042. Gene 461, 5–14 (2010).

40. D. J. Atkinson, M. A. Brimble, Isolation, biological activity, biosynthesis and synthetic studies towards the rubromycin family of natural products. Natural product reports 32, 811–840 (2015).

41. K. B. Herath, H. Jayasuriya, Z. Guan, M. Schulman, C. Ruby, N. Sharma, K. MacNaul, J. G. Menke, S. Kodali, A. Galgoci, Anthrabenzoxocinones from Streptomyces sp. as liver X receptor ligands and antibacterial agents. Journal of natural products 68, 1437–1440 (2005).

42. K. Jiang, X. Yan, Z. Deng, C. Lei, X. Qu, Expanding the Chemical Diversity of Fasamycin Via Genome Mining and Biocatalysis. J Nat Prod 85, 943–950 (2022).

43. W. Yaqing, Y. Quanming, K. James T, N. Lionel M, Generalizing from a few examples: A survey on few-shot learning. ACM computing surveys 53, 1–34 (2020).

44. M. S. Asyaky, R. Mandala, in 2021 8th International Conference on Advanced Informatics: Concepts, Theory and Applications (ICAICTA). (IEEE, 2021), pp. 1–6.

45. K. Lee, K. Lee, H. Lee, J. Shin, A simple unified framework for detecting out-of-distribution samples and adversarial attacks. Advances in neural information processing systems 31, p(2018).

46. K. Ishida, G. Shabuer, S. Schieferdecker, S. J. Pidot, T. P. Stinear, U. Knuepfer, M. Cyrulies, C. Hertweck, Oak-Associated Negativicute Equipped with Ancestral Aromatic Polyketide Synthase Produces Antimycobacterial Dendrubins. Chemistry–A European Journal 26, 13147–13151 (2020).

47. P. Liu, W. Yuan, J. Fu, Z. Jiang, H. Hayashi, G. Neubig, Pre-train, prompt, and predict: A systematic survey of prompting methods in natural language processing. ACM Computing Surveys 55, 1–35 (2023).

48. Y. Tang, J. Weng, P. Zhang, Neural-network solutions to stochastic reaction networks. Nature Machine Intelligence, 1–10 (2023).

49. J. Trinquier, G. Uguzzoni, A. Pagnani, F. Zamponi, M. Weigt, Efficient generative modeling of protein sequences using simple autoregressive models. Nature communications 12, 5800 (2021).

50. L. McInnes, J. Healy, S. Astels, hdbscan: Hierarchical density based clustering. Open Source Softw. 2, 205 (2017).

51. L. McInnes, J. Healy, J. Melville, Umap: Uniform manifold approximation and projection for dimension reduction. arXiv preprint arXiv:.03426, (2018).

52. S. Laine, T. Aila, Temporal Ensembling for Semi-Supervised Learning. arXiv:1610.02242, (2016).

53. S. Andrews. (Babraham Bioinformatics, Babraham Institute, Cambridge, United Kingdom, 2010).

54. R. R. Wick, L. M. Judd, C. L. Gorrie, K. E. Holt, Completing bacterial genome assemblies with multiplex MinION sequencing. Microb Genom 3, e000132 (2017).

55. R. R. Wick, L. M. Judd, C. L. Gorrie, K. E. Holt, Unicycler: Resolving bacterial genome assemblies from short and long sequencing reads. PLoS Comput Biol 13, e1005595 (2017).

56. S. Koren, B. P. Walenz, K. Berlin, J. R. Miller, N. H. Bergman, A. M. Phillippy, Canu: scalable and accurate long-read assembly via adaptive k-mer weighting and repeat separation. Genome Res 27, 722–736 (2017).

57. R. Vaser, I. Sovic, N. Nagarajan, M. Sikic, Fast and accurate de novo genome assembly from long uncorrected reads. Genome Res 27, 737–746 (2017).

58. T. Seemann, Prokka: rapid prokaryotic genome annotation. Bioinformatics 30, 2068–2069 (2014).

59. M. Wang, J. J. Carver, V. V. Phelan, L. M. Sanchez, N. Garg, Y. Peng, D. D. Nguyen, J. Watrous, C. A. Kapono, T. Luzzatto-Knaan, C. Porto, A. Bouslimani, V. Melnik, M. J. Meehan, W. T. Liu, M. Crusemann, P. D. Boudreau, E. Esquenazi, M. Sandoval-Calderon, R. D. Kersten, L. A. Pace, R. A. Quinn, K. R. Duncan, C. C. Hsu, D. J. Floros, R. G. Gavilan, K. Kleigrewe, T. Northen, R. J. Dutton, D. Parrot, E. E. Carlson, B. Aigle, C. F. Michelsen, L. Jelsbak, C. Sohlenkamp, P. Pevzner, A. Edlund, J. McLean, J. Piel, B. T. Murphy, L. Gerwick, C. C. Liaw, Y. L. Yang, H. U. Humpf, M. Maansson, R. A. Keyzers, A. C. Sims, A. R. Johnson, A. M. Sidebottom, B. E. Sedio, A. Klitgaard, C. B. Larson, C. A. B. P D. Torres-Mendoza, D. J. Gonzalez, D. B. Silva, L. M. Marques, D. P. Demarque, E. Pociute, E. C. O’Neill, E. Briand, E. J. N. Helfrich, E. A. Granatosky, E. Glukhov, F. Ryffel, H. Houson, H. Mohimani, J. J. Kharbush, Y. Zeng, J. A. Vorholt, K. L. Kurita, P. Charusanti, K. L. McPhail, K. F. Nielsen, L. Vuong, M. Elfeki, M. F. Traxler, N. Engene, N. Koyama, O. B. Vining, R. Baric, R. R. Silva, S. J. Mascuch, S. Tomasi, S. Jenkins, V. Macherla, T. Hoffman, V. Agarwal, P. G. Williams, J. Dai, R. Neupane, J. Gurr, A. M. C. Rodriguez, A. Lamsa, C. Zhang, K. Dorrestein, B. M. Duggan, J. Almaliti, P. M. Allard, P. Phapale, L. F. Nothias, T. Alexandrov, M. Litaudon, J. L. Wolfender, J. E. Kyle, T. O. Metz, T. Peryea, D. T. Nguyen, D. VanLeer, P. Shinn, A. Jadhav, R. Muller, K. M. Waters, W. Shi, X. Liu, L. Zhang, R. Knight, P. R. Jensen, B. O. Palsson, K. Pogliano, R. G. Linington, M. Gutierrez, N. P. Lopes, W. H. Gerwick, B. S. Moore, P. C. Dorrestein, N. Bandeira, Sharing and community curation of mass spectrometry data with Global Natural Products Social Molecular Networking. Nat Biotechnol 34, 828–837 (2016).

