## Supplementary Information for "Protein language model-based end-to-end type II polyketide prediction without sequence alignment"

### Electrical Supplementary Information

#### Supplementary Figures

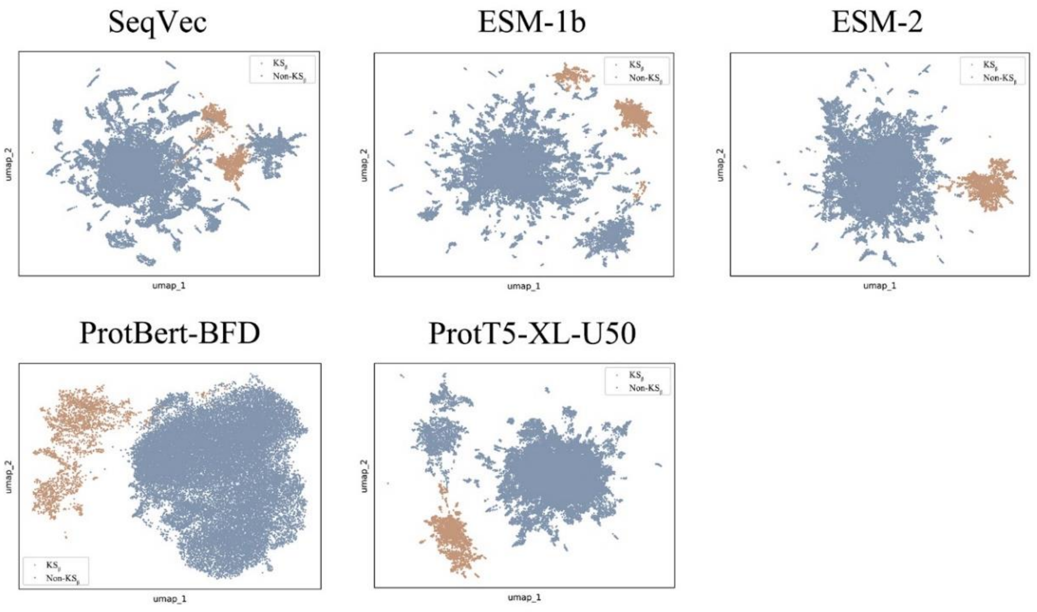


**Fig. S1. Dimensional-reduction representations of $KS_\beta$ and non- $KS_\beta$ embeddings are encoded in five general protein language models.** $KS_\beta$: 2,761; non- $KS_\beta$: Random selection of 36,089 from 761,334.

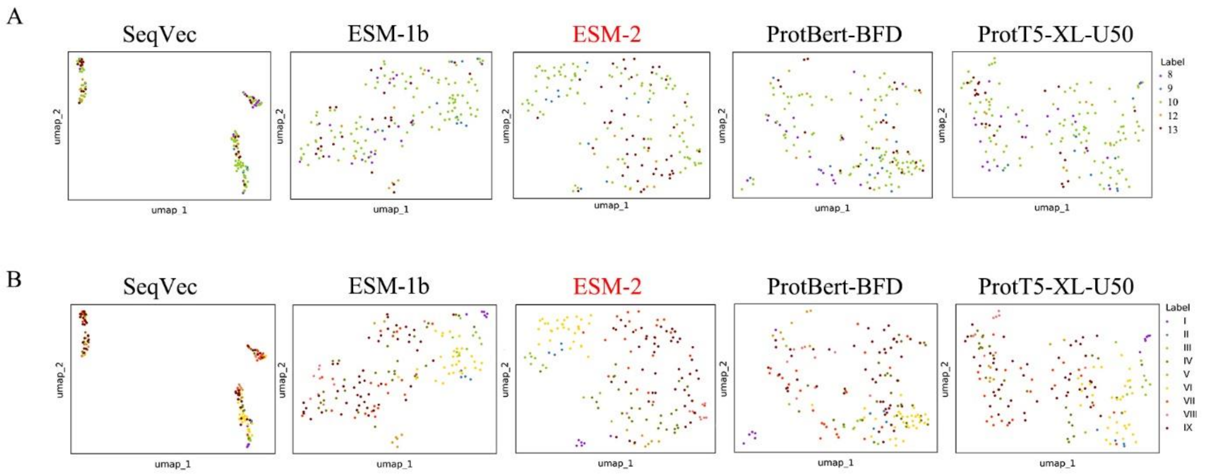


**Fig. S2. Dimensional-reduction representations of 164 $KS_\beta$ embeddings with two types of class label are encoded in five general protein language models.** Panel A: Five class labels according to the building block number of their corresponding to T2PK main skeleton, namely, 8, 9, 10, 12 and 13; Panel B: Nine class labels derived from five class labels using constrained optimization approach. These dimensional-reduction representations were generated through an unsupervised UMAP algorithm.

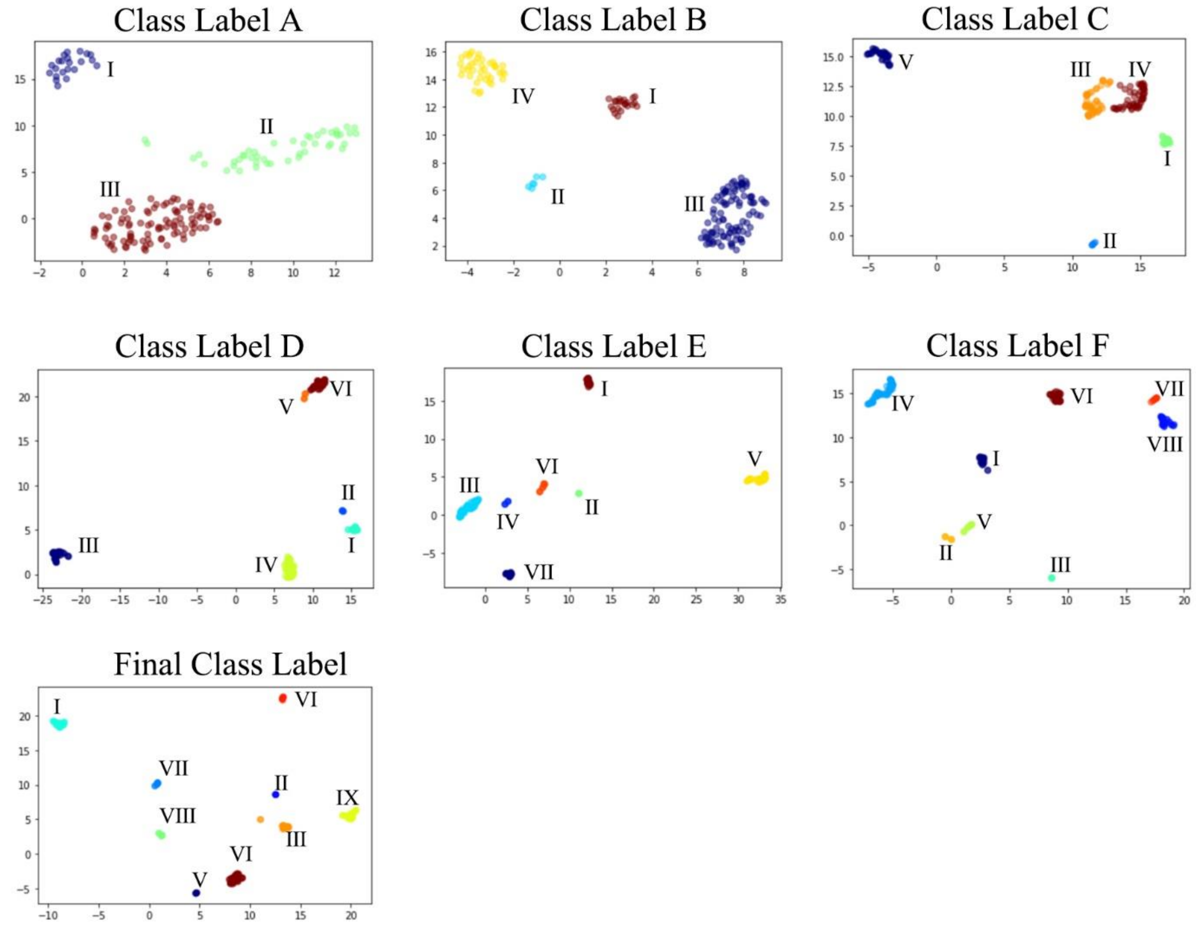


**Fig. S3. Two-dimensional representations showing the process of class labelling for T2PKs.** Details of the hyperparameters used for cluster generation are referred to Table S8.

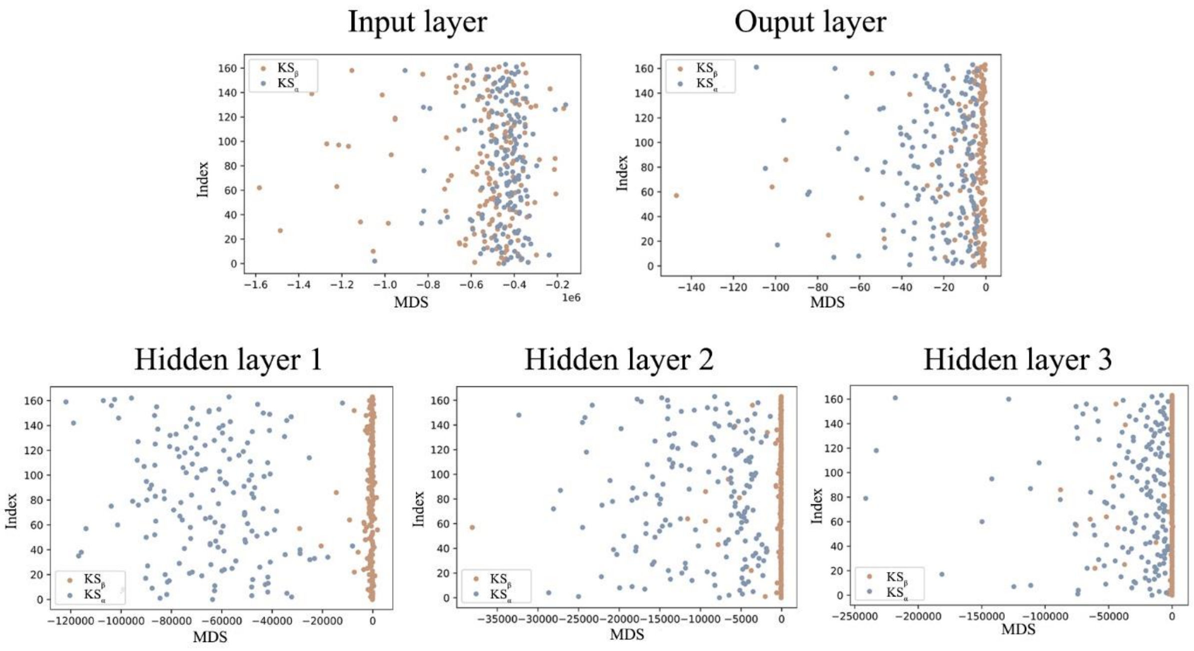


**Fig. S4. Two-dimensional representations of the features extracted from each layer of enhanced T2PK classifier model.** The in-distribution data comprised 164 labelled $KS_{\beta}$, while the out-of-distribution data comprised 164 labelled $KS_{\alpha}$. The Mahalanobis distance-based scores calculated by Gaussian discriminant analysis are plotted on the X-axis, while the Y-axis denotes the index of each datapoint in their dataset.

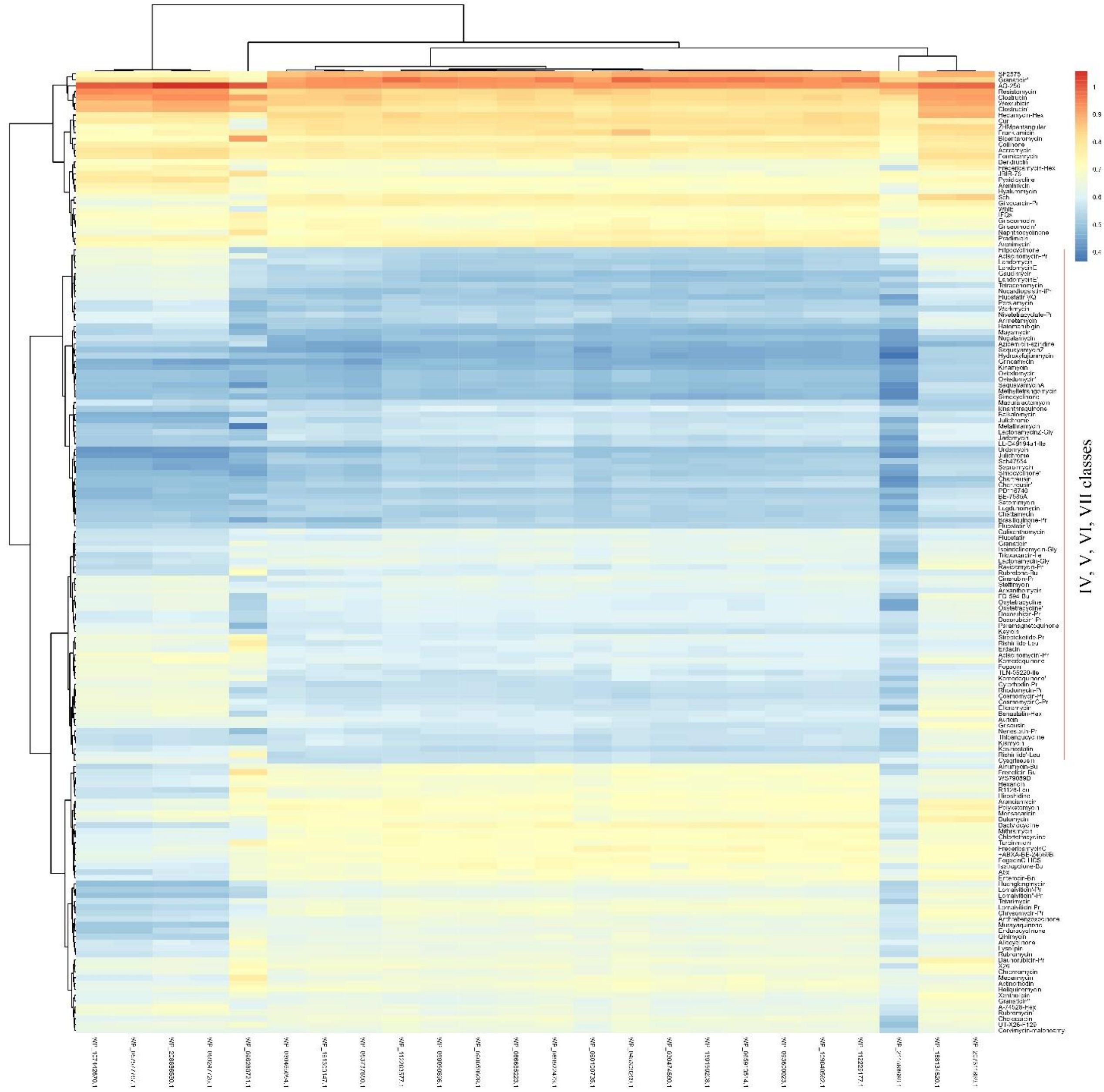


**Fig. S5. Heatmap representation showing the root mean square deviation (RMSD) of the predicted protein structure between 20 novel $KS_\beta$ in ODD cluster 3 and 164 labelled $KS_\beta$.** Detailed values are referred to Table S5.

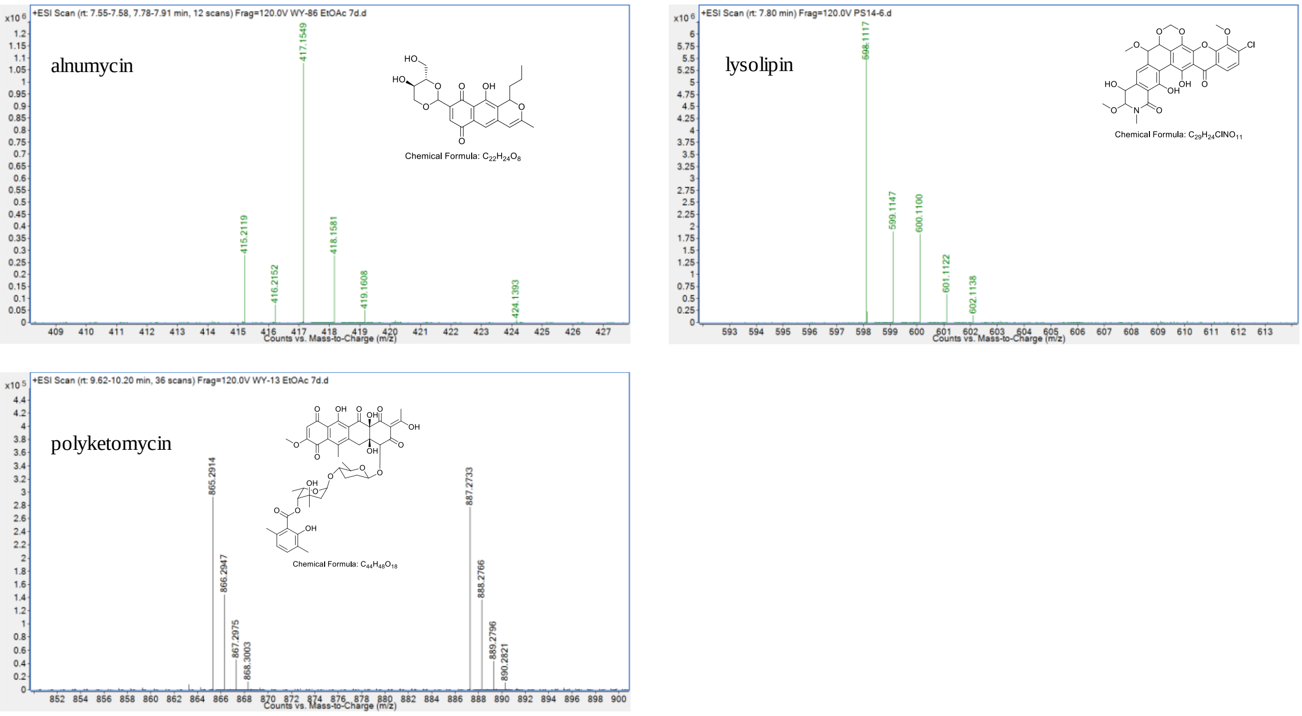


**Fig. S6. Identification of T2PKs detected in this work.** Liquid chromatography high-resolution mass spectrometry (LCHRMS) showing alnumycin (molecular formula $C_{22}H_{24}O_8$, calculated $[M + H]^+$ = 417.1544, observed $[M + H]^+$ = 417.1549, Δ = 1.2 ppm), polyketomycin (molecular formula $C_{44}H_{48}O_{18}$, calculated $[M + H]^+$ = 865.2913, observed $[M + H]^+$ = 865.2914, Δ = 1 ppm), , and lysolipin (molecular formula $C_{29}H_{24}NO_{11}Cl$, calculated $[M + H]^+$ = 598.1111, observed $[M + H]^+$ = 598.1117, Δ = 0.12 ppm), respectively.

**Supplementary Tables**

Note that deo to the scales Table S1-2, S4-5 are deposited as independent Excel spreadsheets.

**Table S3. Performance metrics of $KS_{\beta}$ classifier trained by Random Forest, XGBoost, support vector machine (SVM), and multilayer perceptron (MLP).** TPR: True positive rate; FPR: False positive rate.

| Model | TPR (%) | FPR (%) | Accuracy | Precision | Recall | F1-score |
|---|---|---|---|---|---|---|
| Random Forest | 98.89 | 1.11 | 1.00 | 1.00 | 0.99 | 0.99 |
| XGBoost | 99.63 | 0.37 | 1.00 | 1.00 | 1.00 | 1.00 |
| SVM | 100.00 | 0.00 | 1.00 | 1.00 | 1.00 | 1.00 |
| MLP | 100.00 | 0.00 | 1.00 | 1.00 | 1.00 | 1.00 |

**Table S6. Average root-mean-square deviation (RMSD) calculation between ODD cluster 3 and IV, V, VI, VII classes.**

| | IV | V | VI | VII | ODD cluster 3 |
|---|---|---|---|---|---|
| IV | 0.24 | | | | |
| V | 0.53 | 0.39 | | | |
| VI | 0.48 | 0.57 | 0.29 | | |
| VII | 0.45 | 0.57 | 0.58 | 0.30 | |
| ODD cluster 3 | 0.51 | 0.60 | 0.64 | 0.56 | 0.28 |

**Table S7. Performance comparison on predicting T2PKs using DeepT2, DeepBGC and antiSMASH.**

<table>
<tr><th rowspan="2">Strain name</th><th rowspan="2">Genome Size (bp)</th><th rowspan="2">DeepBGC</th><th colspan="2">Antismash</th><th colspan="4">DeepT2</th></tr>
<tr><th>Counts</th><th>T2 PK name</th><th>Counts</th><th>Class</th><th>T2 PK name</th><th>Euclidean distance</th></tr>
<tr><td rowspan="6">S. sp. PS14</td><td rowspan="6">6848633</td><td rowspan="6">polyketide</td><td rowspan="6">2</td><td rowspan="3">Spore pigment</td><td rowspan="6">2</td><td rowspan="3">VIII</td><td>Cur</td><td>0.72</td></tr>
<tr><td>Sch</td><td>0.98</td></tr>
<tr><td>ZHMpentangular</td><td>1.00</td></tr>
<tr><td rowspan="3">Lysolipin</td><td rowspan="3">IX</td><td>Lysolipin</td><td>0.56</td></tr>
<tr><td>Anthrabenzoxocinone</td><td>0.57</td></tr>
<tr><td>Rubromycin</td><td>0.70</td></tr>
<tr><td rowspan="9">S. sp. WY13</td><td rowspan="9">10713003</td><td rowspan="9">polyketide</td><td rowspan="9">3</td><td rowspan="3">Polyketomycin</td><td rowspan="9">3</td><td rowspan="3">VI</td><td>Dutomycin</td><td>0.13</td></tr>
<tr><td>Polyketomycin</td><td>0.44</td></tr>
<tr><td>Arimetamycin</td><td>0.67</td></tr>
<tr><td rowspan="3">Gaudimycin</td><td rowspan="3">VII</td><td>Gaudimycin</td><td>0.75</td></tr>
<tr><td>Frigocyclinone</td><td>0.85</td></tr>
<tr><td>Landomycin</td><td>0.86</td></tr>
<tr><td rowspan="3">Spore pigment</td><td rowspan="3">VIII</td><td>Cur</td><td>0.65</td></tr>
<tr><td>Collinone</td><td>0.95</td></tr>
<tr><td>ZHMpentangular</td><td>0.97</td></tr>
<tr><td rowspan="3">S. sp. WY86</td><td rowspan="3">11617619</td><td rowspan="3">polyketide</td><td rowspan="3">1</td><td rowspan="3">Alnumycin</td><td rowspan="3">1</td><td rowspan="3">I</td><td>Frenolicin-Bu</td><td>0.39</td></tr>
<tr><td>Alnumycin-Bu</td><td>0.49</td></tr>
<tr><td>Granaticin</td><td>0.81</td></tr>
<tr><td rowspan="9">S. kanamyceticus 4.1441</td><td rowspan="9">10265719</td><td rowspan="9">polyketide</td><td rowspan="9">3</td><td rowspan="3">Aurachin</td><td rowspan="9">3</td><td rowspan="3">I</td><td>Fogacin</td><td>0.37</td></tr>
<tr><td>Isoindolinomycin-Gly</td><td>0.48</td></tr>
<tr><td>Medermycin</td><td>0.58</td></tr>
<tr><td rowspan="3">cinerubin B</td><td rowspan="3">VI</td><td>Keyicin</td><td>0.06</td></tr>
<tr><td>Gilvocarcin-Pr</td><td>0.33</td></tr>
<tr><td>Bipentaromycin</td><td>0.42</td></tr>
<tr><td rowspan="3">Formicamycin</td><td rowspan="3">VIII</td><td>Accramycin(Fasamycin)</td><td>0.74</td></tr>
<tr><td>Formicamycin</td><td>0.79</td></tr>
<tr><td>Sch</td><td>1.20</td></tr>
<tr><td rowspan="3">S. ginsengnesis G7</td><td rowspan="3">9520487</td><td rowspan="3">polyketide</td><td rowspan="3">1</td><td rowspan="3">Spore pigment</td><td rowspan="3">1</td><td rowspan="3">VIII</td><td>Cur</td><td>0.64</td></tr>
<tr><td>Collinone</td><td>0.83</td></tr>
<tr><td>ZHMpentangular</td><td>0.91</td></tr>
<tr><td rowspan="6">S. sp. WY170</td><td rowspan="6">8864212</td><td rowspan="6">polyketide</td><td rowspan="6">1</td><td rowspan="6">Spore pigment</td><td rowspan="6">2</td><td rowspan="3">VI</td><td>Resistomycin</td><td>0.18</td></tr>
<tr><td>Hedamycin-Hex</td><td>0.30</td></tr>
<tr><td>Bipentaromycin</td><td>0.39</td></tr>
<tr><td rowspan="3">VIII</td><td>Collinone</td><td>0.73</td></tr>
<tr><td>Cur</td><td>0.82</td></tr>
<tr><td>Sch</td><td>0.95</td></tr>
</table>

**Table S8. General information on selected hyperparameters to all classifier.**

| Classifier type | Model type | Hyperparameters |
|---|---|---|
| Binary $KS_{\beta}$ classifier | Random forest | n_estimators = 150;<br>max_features='sqrt';<br>min_samples_split=4;<br>min_samples_leaf=1;<br>max_depth=6 |
| | XGBoost | learning_rate=0.1;<br>max_depth=5;<br>min_child_weight=1;<br>subsample=0.7 |
| | Multilayer perceptron | num_epochs=20;<br>learning_rate=0.001;<br>activation='relu';<br>solver='adam';<br>neuron = 50;<br>hidden_layer = 3 |
| | Support vector machine | C=10;<br>gamma=1;<br>kernel=' rbf' |
| Initial T2 PK classifier | Random forest | n_estimators = 150;<br>max_features='log2';<br>min_samples_split=5;<br>min_samples_leaf=1;<br>max_depth=6 |
| | XGBoost | learning_rate=0.2;<br>max_depth=3;<br>min_child_weight=1;<br>subsample=0.6 |
| | Multilayer perceptron | num_epochs=180;<br>learning_rate=0.001;<br>activation='tanh';<br>solver='adam';<br>neuron=200;<br>hidden_layer=3;<br>dropout=0.5<br>train_loader_batch_size=12<br>test_loader_batch_size=3 |
| | Support vector machine | C=10;<br>gamma=1;<br>kernel=' rbf'; |
| Enhanced T2 PK classifier | Multilayer perceptron | num_epochs=500;<br>learning_rate=0.001;<br>activation='tanh';<br>solver='adam';<br>neuron=200;<br>hidden_layer=3;<br>Gaussian_noise_mean=0<br>Gaussian_noise_stddev =0.06<br>train_loader_batch_size=12<br>test_loader_batch_size=3<br>merge_loader_batch_size=64<br><br>Unsupervised loss weight:<br>max_val=50;<br>ramp_up_multi = -2<br>max_epochs=400<br>n_labeled = 164 |

| | | |
|---|---|---|
| | | n_samples = 2761 |

**Table S9. General information on selected hyperparameters for each labelling process.** n_neighbors, n_components, min_dist, random_state are the hyperparameters of UMAP; min_cluster_size, min_samples, cluster_selection_epsilon are the hyperparameters of HDBSCAN; max_evals is the hyperparameter of *Fmin* function from *hyperopt* packages. label_count is the hyperparameter of label cost function. Following hyperparameters were set as default along the tunning**:** n_components = 3; min_cluster_size = 2; min_samples = None; random_state = 42; max_evals = 100.

| **Stage** | **Range for hyperparameters bayesian optimization** | **Optimized hyperparameters** |
|---|---|---|
| Initial Class label to Class label A | n_neighbors = [20, 50], step = 1 | n_neighbors = 46 |
| | min_dist = [0.1, 1], step = 0.1 | min_dist = 1 |
| | cluster_selection_epsilon = [1, 2], step = 0.2 | cluster_selection_epsilon = 1.6 |
| | label_count = 3 | cluster_count = 3 |
| Class label A to Class label B | n_neighbors = [20, 46], step = 1 | n_neighbors = 35 |
| | min_dist = [0.1, 1], step = 0.1 | min_dist = 0.3 |
| | cluster_selection_epsilon = [0.8, 1.6], step = 0.2 | cluster_selection_epsilon = 0.8 |
| | label_count = 4 | cluster_count = 4 |
| Class label B to Class label C | n_neighbors = [20, 35], step = 1 | n_neighbors = 20 |
| | min_dist = [0.02, 0.3], step = 0.01 | min_dist = 0.05 |
| | cluster_selection_epsilon = [0.6, 1.4], step = 0.2 | cluster_selection_epsilon = 0.8 |
| | label_count = 5 | cluster_count = 5 |
| Class label C to Class label D | n_neighbors = [10, 20], step = 1 | n_neighbors = 14 |
| | min_dist = [0.02, 0.1], step = 0.01 | min_dist = 0.02 |
| | cluster_selection_epsilon = [0.2, 1], step = 0.2 | cluster_selection_epsilon = 0.4 |
| | label_count = 6 | cluster_count = 6 |
| Class label D to Class label E | n_neighbors = [5, 14], step = 1 | n_neighbors = 12 |
| | min_dist = [0.02, 0.1], step = 0.01 | min_dist =0.02 |
| | cluster_selection_epsilon = [0.2, 1], step = 0.2 | cluster_selection_epsilon = 0.6 |
| | label_count = 7 | cluster_count = 7 |
| Class label E to Class label F | n_neighbors = [5, 14], step = 1 | n_neighbors = 9 |
| | min_dist = [0.0, 0.05], step = 0.01 | min_dist =0.03 |
| | cluster_selection_epsilon = [0.2, 0.8], step = 0.2 | cluster_selection_epsilon = 0.4 |
| | label_count = 8 | cluster_count = 8 |
| Class label F to Final Class label | n_neighbors = [3, 15], step = 1 | n_neighbors = 7 |
| | min_dist = [0.0, 0.05], step = 0.01 | min_dist = 0.0 |
| | cluster_selection_epsilon = [0.0, 0.8], step = 0.2 | cluster_selection_epsilon = 0.6 |
| | label_count = 9 | cluster_count = 9 |
